# Does pixel/voxel-size limit the measurement of distances in CBCT-tomography?

**DOI:** 10.1101/486167

**Authors:** Deusdedit L. Spavieri, Jonas Bianchi, Jaqueline Ignacio, João R. Gonçalves, Roland Köberle

**Author notes:** Current Address: Braincare SA, São Carlos, São Paulo, Brasil. The work reported here was performed while DLS was associated with **1**.

## Abstract

The necessity to obtain relevant structural information from tomographic images is an all-pervasive step in a host of clinical and research-areas. Cone Beam Computed Tomography (CBCT) is the imaging modality often used among the many available.

Currently approaches to extract structural properties from raw CBCT-images, involve some manual intervention by experts to measure their properties, such as size and displacements of their geometrical structures. Regarding the factors limiting the precision of these measurements, such as voxel-size and image contrast, we find conflicting statements in the literature. It is therefore useful to provide accurate data under well-defined experimental conditions.

Here we present a method and associated software to measure displacements of geometrical structures. We also determined the minimum measureable displacement and minimum detectable defect in terms of voxel size. We select as our geometrical structure a sample of bovine bone and to provide a set of defects, we drilled a pattern of holes into it. We determined the hole’s three-dimensional structures using confocal spectroscopy.

In order to obtain the minimum measurable displacement, we acquired CBCT-tomographies containing a stationary reference and micro-metrically cnc controlled displacements of the sample. We then process these images with our software to extract the distances and compare them with the cnc displacements. All our processing includes a computational interpolation from the voxel-size of 0.35 mm corresponding to our CBCT-tomographies, down to 0.05 mm. We find that sample-displacements can be measured with a precision of ~ 20*μ*, 17 times smaller than the voxel-size of 0.35 mm.

To measure the size of the holes using our CBCT-tomographies, we first register the holes onto a hole-free region of the sample with our software, then overlay the result with the three-dimensional structure obtained from confocal spectroscopy. We find the minimum detectable hole-size to be 0. 7 mm, twice the voxel-size.

## Introduction

The extraction of meaningful geometrical structures from digital images is a fundamental problem in many research areas with innumerable applications, specially in the medical areas [1–3]. Recently we have seen enormous advances in computer-based methods, due to, inter alia: the overall increase of computational power, the development of new algorithms and imaging techniques. The analysis of biomedical images, which has the bonus of immediate practical application, has generated a particularly strong impetus. Here structural quantitative imaging analysis has been used in disease diagnosis, treatment, monitoring and surgical planning. Among the imaging modalities employed in those applications, Cone Beam Computer Tomography (CBCT) has been one of the most frequently used.

CBCT has several advantages, such as: the greater amount of information obtained in comparison with 2D tomographic methods, lower dose delivery to the patient [4,5] in comparison with conventional Computer Tomography [6] and relatively low cost. Yet, in comparison with Computer Tomography, CBCT images are noisier and have lower contrast resolution [7,8].

Almost all of the applications of CBCT images require quantitative analysis of geometric structures, the most important being the measurement their sizes and the measurement of the distance between different structures.

The main limitations in the detection of a particular structure are voxel-size and image-contrast. Periodic patterns with differing spacing have been traditionally used to compare the precision of different setups, with the conclusion that although voxel-size certainly is a limitation to infer the minimum spacing detectable, an accuracy smaller than voxel-size is obtainable [7]. Limitations due to image-contrast have been analyzed recently by [9], who report that up to 30 % false positives can result even for a contrast of 80%.

Furthermore current approaches to extract geometrical properties from raw data usually involve some manual intervention by experts. For example to extract distances from 2- or 3-dimensional CBCT-tomographies a linear measure using a digital caliper is often used to compare the accuracy of different experts [10,11]. With regard to the precision of different types of physical measurements [12–18], we find statements that voxel-size is not important [11,19], whereas a review asserts that the improvement due to better resolution is not well quantified [20]. Yet the size of the defects studied is rather large (≥ 3× voxel-size) and their geometrical characterization lacks precision. Therefore the minimum size of detectable defects is unknown. We conclude that these methods merit further discussions.

In this paper we want to address two issues concerning images acquired by CBCT tomography: *what is the precision with which we are a,ble to*

- *measure distances between displaced, structures*. For this purpose we manufacture a sample of bovine bone, measuring ~ 2*cm* × 3*cm* × 0.5*cm*, shown in Fig.(1C). We actively displace this macroscopic sample and compare its CBCT-tomographies, which also contain a fixed reference, before and after displacement.
- *obtain the minimum size of defects, using CBCT images*. To produce the defects, we drilled a pattern of holes into the sample shown in Fig.(1F). We measured the three-dimensional structure of the holes using confocal spectroscopy [21]. Since the holes have a contrast better than the usually available in real life, as e.g. in the detection of the condylus [9], our sample was immersed in water to approximate these real life conditions.

**Fig 1.**
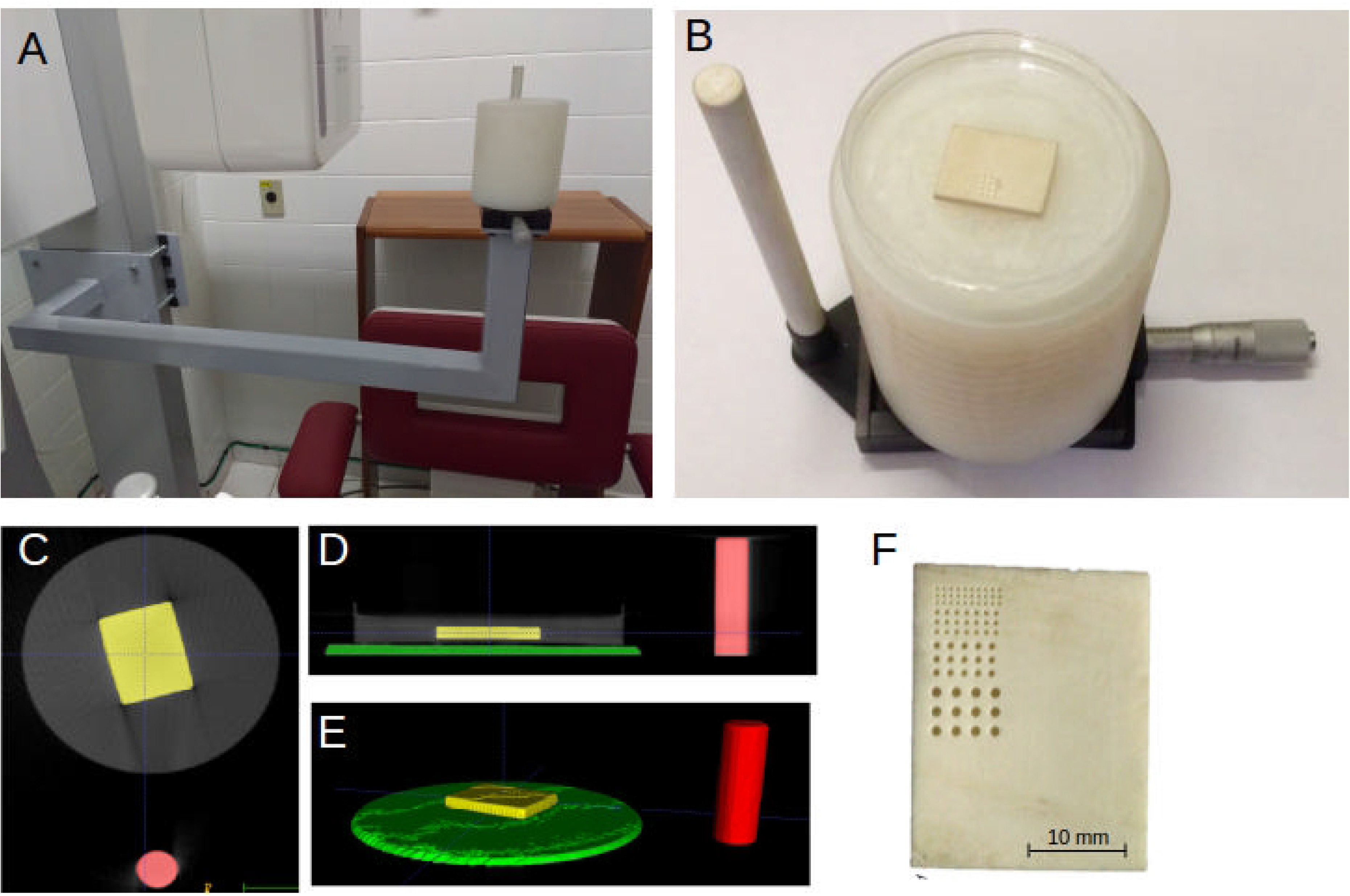
Experimental setup. **A**. Support fixed on the structure of a Scanora 3Dx tomograph. **B**. PVC support to hold the bone sample immersed in water. During the experiment, the sample was moved by the micrometer attached to the sample base, while the vertical PVC rod on the left provided a fixed reference. **C**. Vertical view of the support tomography. Bone sample in yellow and reference rod in red. **D**. Horizontal view of the support tomography. Bone sample in yellow, reference rod in red and PVC base in green. **E**. Three-dimensionally rendered view of the semi-automatically segmented image. **F**. Details of sample, showing the holes with diameters of 0.35,0.5, 0.7 and 1.0 mm.

To obtain the minimum detectable displacement we use the setup shown in Fig.(1A). We accurately displace the sample using a micrometer with 10*μ* precision. We obtain CBCT-tomographies (voxel-size = 0.35 mm) of the whole setup shown in Fig.(1B) and extract the region corresponding to the bone-sample before and after displacements. We describe the registration algorithm used to register the sample.

To obtain the minimum size of defects discernible with our algorithm, we first register the holes onto a hole-free region of the sample using our algorithm. The registered images (color-coded) together with the hole’s profile obtained by confocal spectroscopy (shown in black) are then overlaid with the result shown in Fig(3).

## Methods

### Sample preparation

For data-acquisition we prepared a sample with rectangular cross-section of 2*cm* × 3*cm* and a depth of 0.5*cm* from a piece of bovine tibia-cortex bone, sectioned with a diamond-shaped cutting machine (ISOMET 1000, Buheler IL, USA). The anatomical structure of this sample should be typical for many real life situations and we therefore expect our results to offer a useful benchmark for many applications. We then polished the sample under constant irrigation with a polishing machine (Buehler, Lake Bluff, IL, USA), using a sequence of silicon carbide granules (150,320,400,600,1200,1600,3000) and finished it with diamond paste and felts.

We drilled several holes of different diameters and depths on the surface of the sample (Fig1F) using a computer-numerically-controlled (cnc) mill. The diameter *D_H_* of the holes, their number *N_H_* and mutual distance between holes *d*(*mm*) are shown in the Table 1. The axis of the holes will be called *Z*-axis, which points vertically.

**Table 1.** Sample properties - diameters, numbers of holes and their mutual distances.

| $D_H$ [mm] | $N_H$ | $d$ [mm] |
| --- | --- | --- |
| 0.35 | 30 | 0.6 |
| 0.5 | 21 | 0.7 |
| 0.7 | 15 | 0.8 |
| 1.0 | 15 | 1.2 |

### Hole characterization

In order to infer the size of the smallest holes, which we can detect using tomographic images processed by our algorithm, we need a precise enough measure of their spatial geometry. We therefore used confocal microscopy [21] (Lext 3D Laser Measuring, Center Valley, PA, USA) using laser wavelength of 405*nm* to obtain the three-dimensional structure of the holes. This technique uses a laser to produce fluorescent emission of the sample’s tissue. The fluorescent light is detected only very close to the focal plane. This allows an extremely accurate spatial resolution - 0.001 mm - of the sample’s three-dimensional geometry.

### Translation Stage and Image acquisition

We used a Scanora 3Dx (Soredex, Finland) tomograph with a field-of-view (FOV) measuring 165 × 80*mm*, with voxel size of 0.35 mm at tension 90 kV, current 10 mA and exposition time of 2.4 seconds. Again as mentioned in the section *Sample preparation* this setup may be considered typical for many application. Of course the details of our conclusions reached, when using another tomograph, will depend on experimental parameters.

On the tomograph we attached a steel arm supporting a translation stage (PT1, Thor Labs) equipped with a movable basis controlled by a micrometer (Mitutoyo) of 10 *μ* per division. We fixed a vertical PVC rod on the static part of the translation stage providing a reference to measure distances from. We mounted a PVC support to hold a Petri-dish containing the sample (Fig 1 A, B) on the movable part of the translation stage. The dish was filled with water, just enough to cover the sample, in order to simulate the contrast of the bone surface equal to natural conditions found in living tissues.

To measure the minimum displacement detectable by the our registration algorithm, we took 11 CBCT-tomographies of the sample, moved by a step of 0.1 mm in the Y-direction after each take. Here the (*X, Y*)-directions are two orthogonal axes in a horizontal plane.

### Measuring Image Displacement/Distances

We aim to provide a benchmark for the precision with which distances between well defined geometrical structures can be measured. For this purpose we select as geometrical structure our macroscopic sample (with cross-section of 2*cm* × 3*cm*) shown in Fig.(1 C), whose displacement is to be measured. The structure to be displaced, shown in Fig.(1 B) contains not only our sample, but also the dish filled with water and the vertical PVC rod, which acts as a static reference.

We segment the volumetric image of the whole setup Fig.(1 B), using the *region competition algorithm* [22] implemented in the ITK-SNAP segmentation software [23], before and after displacement to extract the regions of interest: *bone sample, circular dish and rod*. Yet the borders of the bone sample, which will be used to measure displacements, are not always well defined. To refine their positions, we expand the region labeled *bone sample* via a level set contour propagation [24] followed by an AND operation between the *bone sample* region before and after expansion. A binary threshold filter [25] is then sufficient to obtain a precisely labeled *bone sample*. This eliminates eventual water regions adjacent to the *bone sample*.

We first increased the image-resolution by a factor of seven reducing the voxel-size from 0.35 mm to 0.05 mm, using cubic spline interpolation [26]. We then registered only the regions corresponding to the *bone sample* to compute the *Distance Transform* [27] between the *original* and the *displaced* images. We define the displacement to be the difference between the centroids of the original and the transformed *bone sample* regio The pseudo-code^1^ for these steps runs as follows:

#### Algorithm 1

Displacement calculation

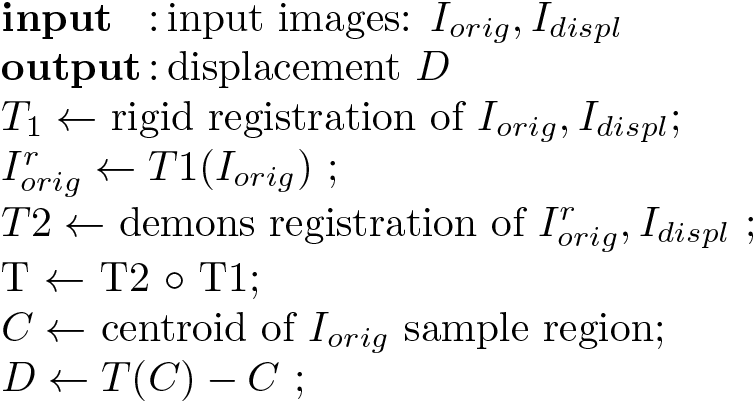

### Hole detection

In order to measure the size of the minimal hole detectable with our registration process, we again increased the resolution sevenfold to a voxel size of 0.05 mm and applied a curvature anisotropic diffusion filter [26,28] to reduce image noise. We then registered a region of the sample without holes onto the region with holes using the demons algorithm [29]

We then overlayed the registered images with the hole’s profile obtained by confocal spectroscopy as follows. Given a hole’s image, we measured the coordinates of the hole’s center. Out of the 3-dimensional image we cut a plane containing the center and overlayed the plane with the hole’s profile such that the centers coincide. We resized the plane’s image such that its scale coincides with the scale used in the spectroscopy profile. The result is shown in Fig(3)

## Results and discussion

### Minimum measurable displacement

To ascertain the minimum detectable displacement, in particular whether it could be less than the voxel size, we plotted the displacements *D_m_* of the 11 CBCT-images vs. the *nominal* micro-metrically controlled ones *D_μ_* in Fig.(2)A. The *X, Z*-displacements are small, but not exactly zero, meaning that the sample did not move exactly in the *Y*-direction. We plotted the distributions of the *Differential displacement D_m_* − *D_μ_* in Fig.(2)B. Their standard-deviations in the *X, Y, Z*-directions are ~ 9.2, 2, 0.4 *μ* respectively. We attributed the somewhat larger *X*-deviation in part to slower convergence of the software-algorithm for this direction. Since the micrometer precision is ~ 20*μ* and we estimated errors in the geometry also to be of this size, we conclude that the precision of the displacement-measurement is limited to 20*μ*, ~ 17 times smaller than the original voxel-size of 350*μ*.

**Fig 2.**
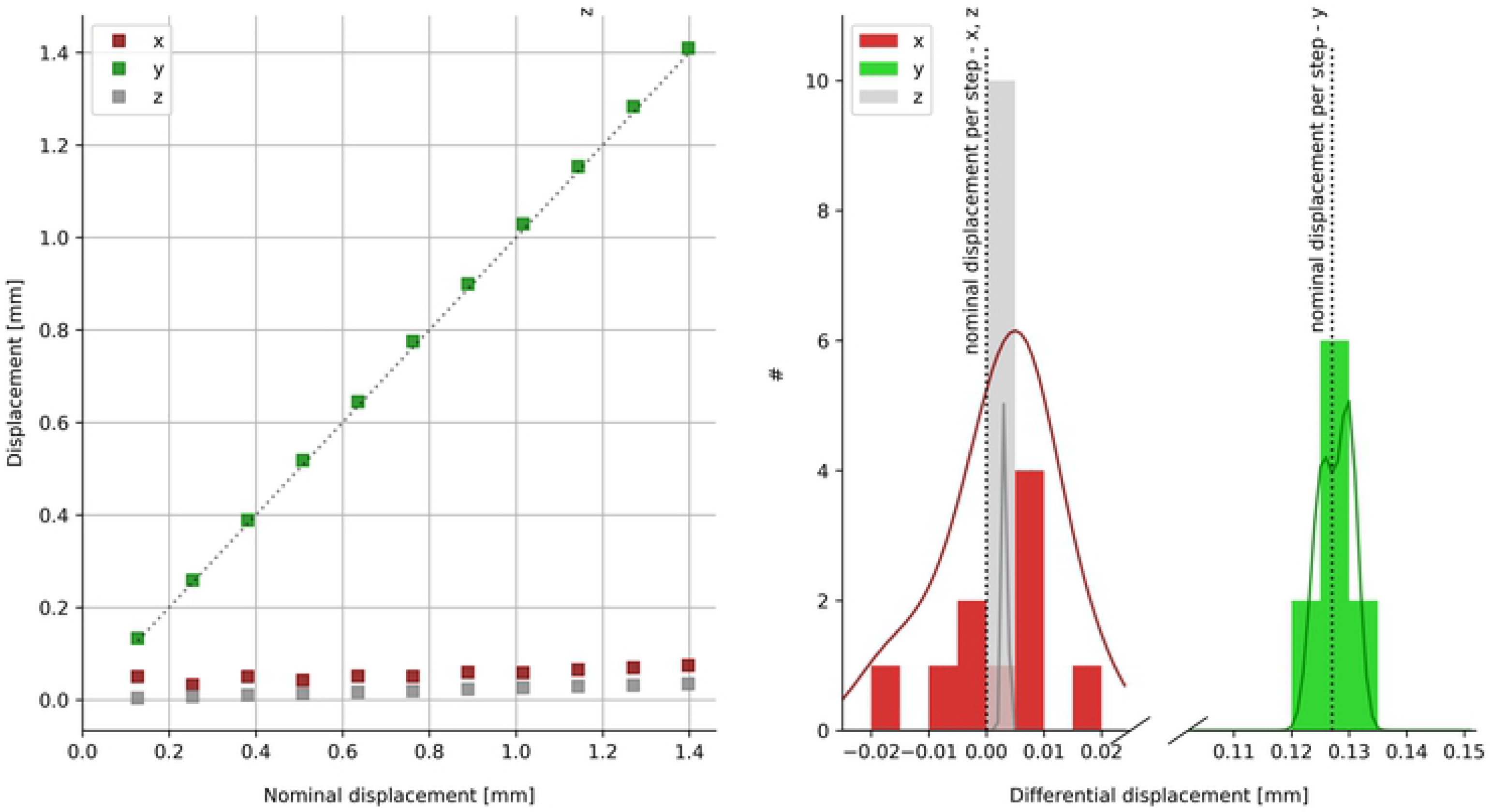
Displacements (Voxel size of 0.35 mm). **A**. Measured *D_m_* vs. micro-metrically controlled *D_μ_* displacements. **B**. Distribution of differential displacements *D_m_* − *D_μ_* in each direction between steps. Continuous lines represent a smoothed continuous version of the discreet differential displacement’s histograms using kernel density estimations.

### Minimum detectable hole size

To estimate the minimum size of the detectable holes, we used the images of the holes, which have been registered by our algorithm onto the hole-free bone sample. We then compared these registered images (color-coded) with the hole’s profile obtained by confocal spectroscopy (in black) in Fig(3).

**Fig 3.**
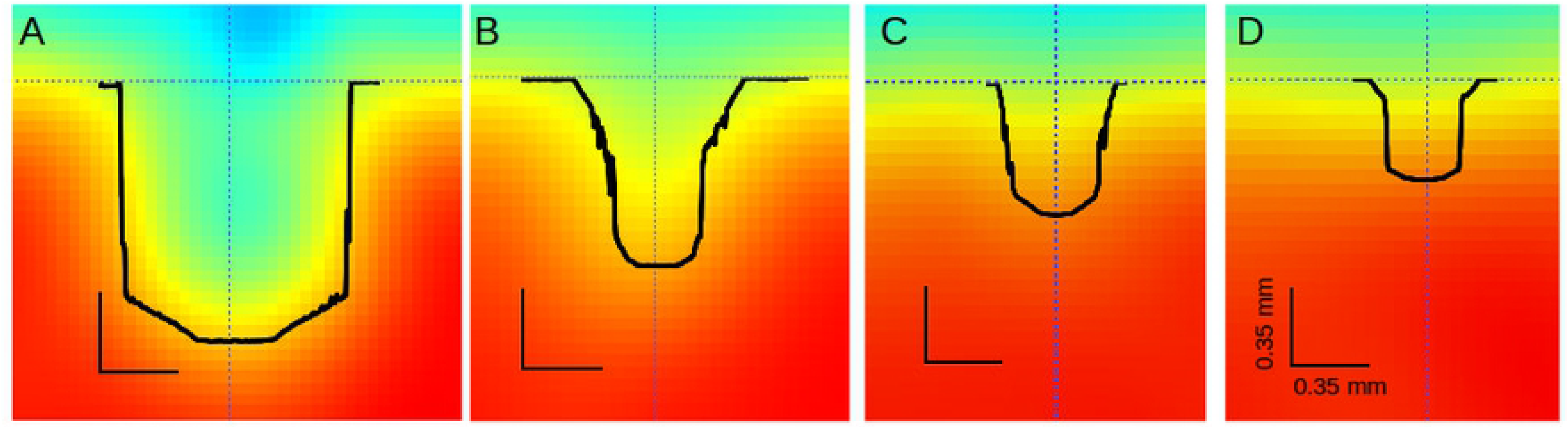
The 3-dimensional shape of the holes listed in Table I, mapped out by confocal spectroscopy - shown in black. The results of our software analysis, which used the pixel-size of 0.35 * 0.35mm shown by the black right-angles, is color-coded. **A**: *D_H_* = 1*mm* **B**: *D_H_* = 0.7*mm* **C**: *D_H_* = 0.5*mm* **D**: *D_H_* = 0.35.*mm*.

We see that holes of sizes ≥ 0.7 mm - Fig(3 A,B) - are very well detected, but not the smaller holes. We note however, that the limiting factor here is not the detection-algorithm, but the voxel-size, as already noted by [20], since the DICE coefficient [30] turns out to be 0.99. Two experts with several years experience using the semi-automatic ITK-SNAP software [23] to segment the image were unable to label even the 0.5mm diameter holes.

## Conclusion

Using our software algorithm, based on open source code, we showed how to obtain

1. the precision with which the distance between displaced bone structures can be measured from their acquired CBCT images;
2. the minimum detectable size of bone defects using those images.

In our displacement measurements we used a bovine bone sample with cross-section of 2*cm* × 3*cm* and a depth of 0.5*cm*. The anatomical structure of this sample should be typical for many real life situations and we therefore expect our results to offer a useful benchmark for many applications. We take special care to precisely extract the sample’s borders, since their displacements are to be measured. Computationally increasing the image-resolution seven times, we are able to measure displacements with a precision of 20*μ*, which is ~ 17 times smaller than the voxel-size of 0.35 mm. We also obtain a minimum detectable defect-size of ≥ 0.7 mm, which was twice the voxel-size.

Although we used data acquired with one tomograph only, with images acquired using just one set of configurations, we expect our results to be very robust relative to the voxel-size.

## Acknowledgments

The confocal measurements were executed by Dr. Alex C. Bottene, member of Prof. Reginaldo T. Coelho’s group, Laboratory of Advanced Processes and Sustainability - LAPRAS, School of Engineering at São Carlos, University of São Paulo. We thank G. F. Brito de Almeida, member of Prof. Cleber Mendonca de Barros’s group at the Instituto de Física, for exploring the geometry of the bone-landmarks with perfilometry methods. JRG was partially funded by Fundação de Amparo a Pesquisa do Estado de São Paulo (FAPESP)-contract 2013/05031-8. JB was partially funded by Coordenação de Aperfeicoamento de Pessoal de Nivel Superior (CAPES)-Demanda Social. It is our pleasure to acknowledge the excellent work executed by the machine shop at the Instituto de Física de São Carlos.

## Footnotes

1 The demons registration step *T*2 in the algorithm is actually the identity here, since the original and the displaced sources for the CBCT-images are identical. It is included, because it is part of our software package, which aims at comparing differently distorted sources.

## References

1. Ashton Ea. Quantitative Medical Imaging. Journal of Imaging Science and Technology. 2007;51(2):117. doi:10.2352/J.ImagingSci.Technol.(2007)51:2(117).

2. Kim TY, Son J, Kim KG. The recent progress in quantitative medical image analysis for computer aided diagnosis systems. Healthcare Informatics Research. 2013;17(3):143–149. doi:10.4258/hir.2011.17.3.143.

3. Suetens P. Fundamentals of medical imaging. Cambridge university press; 2017.

4. Patel S. New dimensions in endodontic imaging: Part 2. Cone beam computed tomography. International Endodontic Journal. 2009;42(6):463–475. doi:10.1111/j.1365-2591.2008.01531.x.

5. Kapila SD, Nervina JM. CBCT in orthodontics: assessment of treatment outcomes and indications for its use. Dento maxillo facial radiology. 2014;44(1):20140282. doi:10.1259/dmfr.20140282.

6. Nicolielo LFP, Van Dessel J, Shaheen E, Letelier C, Codari M, Politis C, et al. Validation of a novel imaging approach using multi-slice CT and cone-beam CT to follow-up on condylar remodeling after bimaxillary surgery. International Journal of Oral Science. 2017;9(3):139–144.

7. Brüllmann D, Schulze RKW. Spatial resolution in CBCT machines for dental/maxillofacial applications - What do we know today? Dentomaxillofacial Radiology. 2015;44(1). doi:10.1259/dmfr.20140204.

8. Lechuga L, Weidlich GA. Cone Beam CT vs. Fan Beam CT: A Comparison of Image Quality and Dose Delivered Between Two Differing CT Imaging Modalities. Cureus. 2016;8(9).

9. Jones EM, Papio M, Tee BC, Beck FM, Fields HW, Sun Z. Comparison of cone-beam computed tomography with multislice computed tomography in detection of small osseous condylar defects. American Journal of Orthodontics and Dentofacial Orthopedics. 2016;150(1):130–139. doi:10.1016/j.ajodo.2015.12.019.

10. Kamburoglu K, Kolsuz E, Kurt H, Kiliç C, Ozen T, Paksoy CS. Accuracy of CBCT measurements of a human skull. Journal of Digital Imaging. 2011;24(5):787–793. doi:10.1007/s10278-010-9339-9.

11. Güngör E, Doğan MS. Reliability and accuracy of cone-beam computed tomography voxel density and linear distance measurement at different voxel sizes: A study on sheep head cadaver. Journal of Dental Sciences. 2017;12(2):145–150. doi:10.1016/j.jds.2016.11.004.

12. Repesa M, Sofic A, Jakupovic S, Tosum S, Kazazic L, Dervisevic A. Comparison of results of measurement of dimensions of the placed dental implants on cone beam computed tomography with dimensions of the producers of the implants. Acta Informatica Medica. 2017;25(2):116–120. doi:10.5455/aim.2017.25.116-120.

13. Timock AM, Cook V, McDonald T, Leo MC, Crowe J, Benninger BL, et al. Accuracy and reliability of buccal bone height and thickness measurements from cone-beam computed tomography imaging. American Journal of Orthodontics and Dentofacial Orthopedics. 2011;140(5):734–744. doi:10.1016/j.ajodo.2011.06.021.

14. Stratemann S, Huang J, Maki K, Miller A, Hatcher D. Comparison of cone beam computed tomography imaging with physical measures. Dentomaxillofacial Radiology. 2008;37(2):80–93. doi:10.1259/dmfr/31349994.

15. Sun L, Zhang L, Shen G, Wang B, Fang B. Accuracy of cone-beam computed tomography in detecting alveolar bone dehiscences and fenestrations. American Journal of Orthodontics and Dentofacial Orthopedics. 2015;147(3):313–323. doi:10.1016/j.ajodo.2014.10.032.

16. Baumgaertel S, Palomo JM, Palomo L, Hans MG. Reliability and accuracy of cone-beam computed tomography dental measurements. American Journal of Orthodontics and Dentofacial Orthopedics. 2009;136(1):19—25. doi:10.1016/j.ajodo.2007.09.016.

17. Berco M, Rigali PH, Miner RM, DeLuca S, Anderson NK, Will LA. Accuracy and reliability of linear cephalometric measurements from cone-beam computed tomography scans of a dry human skull. American Journal of Orthodontics and Dentofacial Orthopedics. 2009;136(1):17.e1–17.e9. doi:10.1016/j.ajodo.2008.08.021.

18. Damstra J, Fourie Z, Huddleston Slater JJR, Ren Y. Accuracy of linear measurements from cone-beam computed tomography-derived surface models of different voxel sizes. American Journal of Orthodontics and Dentofacial Orthopedics. 2010;137(1):16.e1–16.e6. doi:10.1016/j.ajodo.2009.06.016.

19. Ganguly R, Ramesh A, Pagni S. The accuracy of linear measurements of maxillary and mandibular edentulous sites in conebeam computed tomography images with different fields of view and voxel sizes under simulated clinical conditions. Imaging Science in Dentistry. 2016;46(2):93–101. doi:10.5624/isd.2016.46.2.93.

20. Spin-Neto R, Gotfredsen E, Wenzel A. Impact of voxel size variation on CBCT-based diagnostic outcome in dentistry: A systematic review. Journal of Digital Imaging. 2013;26(4):813–820. doi:10.1007/s10278-012-9562-7.

21. Pawley J. Handbook of Biological Confocal Microscopy. third edition ed. Berlin: Springer; 2006.

22. Zhu SC, Yuille A. Region competition: Unifying snakes, region growing, and Bayes/MDL for multiband image segmentation. Pattern Analysis and Machine Intelligence. 1996;18(9).

23. Yushkevich PA, Piven J, Hazlett HC, Smith RG, Ho S, Gee JC, et al. User-guided 3D active contour segmentation of anatomical structures: significantly improved efficiency and reliability. NeuroImage. 2006;31(3):1116–1128. doi:10.1016/j.neuroimage.2006.01.015.

24. Sethian J. Level Set Methods and Fast Marching Methods. Cambridge University Press; 1999.

25. Ibanez L, Schroeder W, Ng L, Cates J. The ITK Software Guide.. vol. 2. ITKSoftware; 2005; 2005. Available from: http://www.itk.org/ItkSoftwareGuide.pdf

26. Johnson HJ, McCormick M, Ibáñez L, Consortium TIS. The ITK Software Guide; 2013. Available from: http://www.itk.org/ItkSoftwareGuide.pdf.

27. Danielsson PE. Euclidean distance mapping. Computer Graphics and image processing. 1980;14(3):227–248.

28. Perona P, Shiota T, Malik J. Anisotropic diffusion. In: Geometry-driven diffusion in computer vision. Springer; 1994. p. 73–92.

29. Vercauteren T, Pennec X, Perchant A, Ayache N. Diffeomorphic demons: efficient non-parametric image registration. NeuroImage. 2009;45(1 Suppl):S61–72. doi:10.1016/j.neuroimage.2008.10.040.

30. Zou KH, Warfield SK, Bharatha A, Tempany CMC, Kaus MR, Haker SJ, et al. Statistical validation of image segmentation quality based on a spatial overlap index. Academic radiology. 2004;11(2):178–189. doi:10.1016/S1076-6332(03)00671-8.

